# Expression kinetics and innate immune response after electroporation and lipid nanoparticle mediated delivery of a self-amplifying mRNA in the skin of mice

**DOI:** 10.1101/528612

**Authors:** Hanne Huysmans, Zifu Zhong, Joyca De Temmerman, Barbara L. Mui, Ying K. Tam, Séan Mc Cafferty, Arlieke Gitsels, Daisy Vanrompay, Niek N. Sanders

## Abstract

In this work we studied the expression kinetics and innate immune response of a self-amplifying mRNA (sa-RNA) after electroporation and lipid nanoparticle (LNP) mediated delivery in the skin of mice. Intradermal electroporation of the sa-RNA resulted in a plateau-shaped expression with the plateau between day 3 and 10. The overall protein expression of sa-RNA was significant higher than that obtained after electroporation of pDNA or non-replication mRNAs. Moreover, intradermal electroporation of sa-RNA induced a short-lived innate immune response that did not affect the expression of the sa-RNA. A complete different expression profile and innate immune response was observed when LNPs were used. The expression peaked 24h after intradermal injection of sa-RNA-LNPs and subsequently showed a sharp drop. This drop can be explained by the strong innate immune response elicited by the sa-RNA-LNPs 4h after injection. Interestingly, sa-RNA-LNPs were able to transfection the draining lymph nodes after intradermal injection.

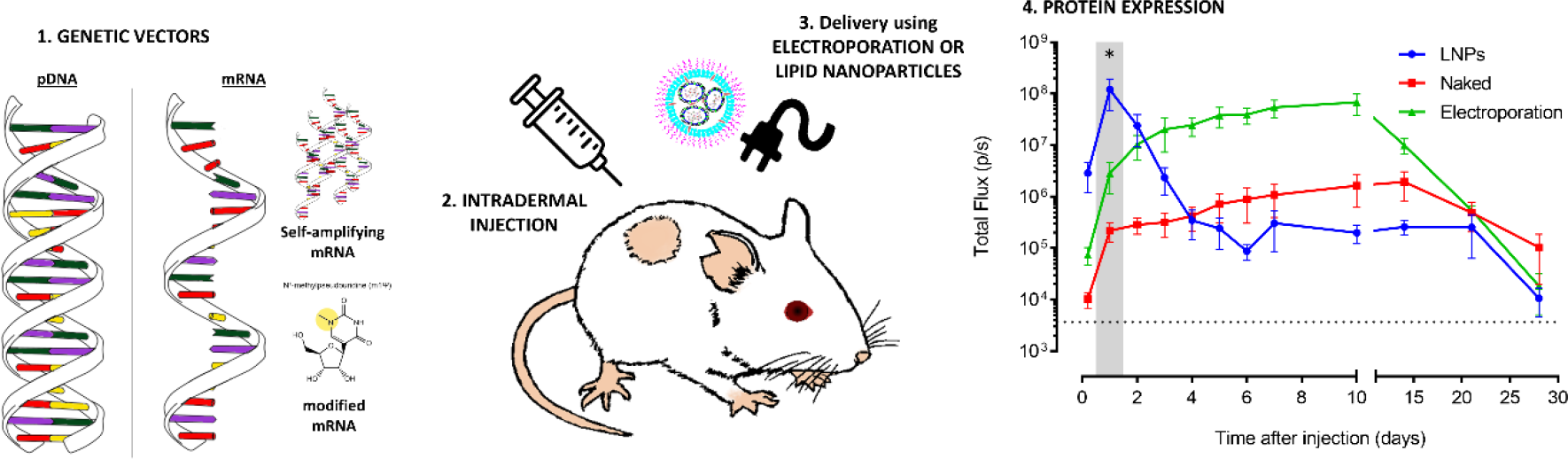

## 1. Background

Synthetic mRNAs are currently intensively studied for protein (replacement) therapy, gene editing, stem cell reprogramming and immunotherapy [1–3]. Each application requires distinct mRNA properties. For instance, protein (replacement) therapy asks for a long-acting and innate immunosilent mRNA, while mRNA vaccination may require opposite characteristics. Multiple synthetic mRNA platforms, such as unmodified mRNA, nucleoside-modified mRNA and self-amplifying mRNA (sa-RNA) are currently available. To find the right match between the foreseen therapeutic application and the mRNA platform, information on the *in vivo* expression kinetics and innate immune response of these different mRNA platforms is crucial. Nucleoside-modified mRNAs are considered innate immunosilent [4], and sa-RNAs are long-acting as they contain the coding sequences of a viral replicase complex that ensures amplification of the complete sa-RNA strand and especially the shorter subgenomic mRNA strand that contains the gene of interest [5–7]. The amplification and abundance of these subgenomic mRNAs engenders a high protein production. The viral replicase complex in sa-RNA originates from a single-stranded positive-sense RNA virus like e.g. Venezuelan equine encephalitis virus (VEEV) [7–10].

For the *in vivo* delivery of synthetic mRNA non-viral carriers, like lipid nanoparticles (LNPs) [11, 12] (reviewed in [13]), as well as physical methods, like electroporation [14–16], have been used. Non-viral carriers formulate the mRNA into nanoparticles that enter the cells by endocytosis, while it is believed that physical methods like electroporation mainly deliver the mRNA directly in the cytosol via temporal cell membrane perforations [12, 17]. One can expect that these different delivery mechanisms will influence the expression efficacy as well as the extent to which the innate immune system recognizes the synthetic mRNA. The recognition of synthetic mRNA by the innate immune system occurs by pathogen recognition receptors (PRRs) that are located in the endosomes (Toll like receptor (TLR) 3, 7 and 8) and in the cytoplasm (e.g. retinoic acid-inducible gene I- (RIG-I-) like receptors). Activation of these PRRs results in the induction of NF-κB, caspase 1, and interferon regulatory factor 3 and 7 (IRF3 and IRF7), which respectively leads to the production of pro-inflammatory cytokines, cell death and type I interferons [18]. Type I interferons are known to activate 2’,5’-oligoadenylate synthetase (OAS) and protein kinase R (PKR) [10]. The former subsequently activates an endonuclease (RNaseL) that degrades the intracellular mRNA, while the latter inhibits the translation by phosphorylating eIF2 [19, 20]. These cellular actions will decrease the translation of the introduced synthetic mRNA. Furthermore, extensive activation of caspase-1 by synthetic mRNA can also cause pyroptosis, a form of immunogenic cell death that exhibits features of apoptosis as well of necrosis [18, 21]. For certain applications like protein (replacement) therapy a repeated administration of the mRNA will be required. For these applications it will not only be important to control the extent of the innate immune response but also the duration of the innate immune response. Indeed, if the innate immune response has not faded away at the moment of the next mRNA injection a chronical inflammatory condition will be established. The duration of the innate immune response after *in vivo* delivery of mRNA is not well studied and most information comes from studies that used RT-PCR or ELISA to quantify key innate immune proteins [22–24]. Although RT-PCR is a very sensitive technique, the correlation between mRNA levels and protein expression are not always linear [25]. On the other hand, ELISA may not be sensitive enough to detect moderate innate immune responses in the excised tissues [26, 27]. Furthermore, analyzing the *in vivo* innate immune response in tissues by RT-PCR and ELISA is invasive and therefore cannot be used to follow-up the innate immune response in one and the same mouse. Therefore, reporter mice like e.g. the IFN-β luciferase reporter mice may serve as a sensitive and reliable alternative to monitor the extent, duration and location of the innate immune responses after mRNA delivery in mice [27].

Recently, we developed, in collaboration with the laboratory of Ron Weiss (Department of Biological Engineering, Massachusetts Institute of Technology, USA), a new sa-RNA. This sa-RNA has the capacity to overcome innate immune responses as it encodes the non-structural proteins (nsPs) of VEEV, which are known to impair critical signaling events downstream of the type I IFN receptor, leading to disruption of STAT1 signaling [28]. In this study we determined the *in vivo* efficacy and innate immunity of this novel VEEV-based sa-RNA at different doses and benchmarked it against pDNA, unmodified and m1ψ-modified synthetic mRNA. To avoid carrier related effects on the efficiency and innate immunogenicity of these vectors, *in vivo* electroporation was used to compare the *in vivo* performance of these vectors. Electroporation and carrier mediated delivery of mRNA occurs by completely different mechanism. As it is currently not known to which extend these different delivery mechanisms affect the *in vivo* performance of synthetic mRNA, we formulated our sa-RNA into LNPs and compared the expression profile and innate immune response with that obtained after *in vivo* electroporation. Because of our interest in mRNA vaccination, intradermal delivery was used as the skin is extremely immune-competent and easily accessible [29]. Our interest in mRNA vaccination also inspired us to study the mRNA expression in the draining lymph nodes as an indicator of mRNA transfection in antigen presenting cells.

## 2. Methods

### 2.1 Mice

Female wild-type BALB/cJRj mice were purchased from Janvier (France) and housed in individual ventilated cages in a climate-controlled facility under a 14/10h light/dark cycle. Heterozygous interferon-β reporter mice with a BALB/c background were a kind gift of Johan Grooten [30] and the breed was further maintained in-house. All mice were aged between 7-10 weeks at the start of the experiments and kept in individually ventilated cages with ad libitum access to feed and water. The *in vivo* mice experiments were conducted with the approval of the Ethical committee of the Faculty of Veterinary Medicine, Ghent University (EC2016/17).

### 2.2 Plasmid constructs

Bacteria containing the pGL4.13 plasmid (GenBank: AY738225) were a kind gift of Katrien Remaut (Ghent University). This plasmid from Promega (Wisconsin, USA) encodes the reporter gene luciferase (*luc2)* from *P. pyralis* controlled by the SV40 promoter and was used to examine expression kinetics. Similarly, a pDNA (pEGFP-N1, GenBank: U55762.1) encoding eGFP was used to study the IFN-β response after intradermal electroporation of pDNA in reporter mice. *E. coli* containing the pEGFP-N1 were also a kind gift of Katrien Remaut. This plasmid contains the eGFP gene controlled by a CMV promoter.

Further, we used 4 different plasmids for the production of RNA by *in vitro* transcription (IVT). PTK160 (11519 base pairs (bp)) and pMC15 (10586bp) are derived from VEEV strain TC-83 containing a substitution in the 5’UTR (r.3a>g) and in nsP2 (p.Q739L). The sequence coding for the structural proteins was replaced by the reporter gene luc2 or eGFP respectively, corresponding to the sequences used in pGL4.13 and pEGFP-N1. Additionally, the vectors contained restriction sites for the I-SceI endonuclease prior to a T7 polymerase promotor and downstream of a short poly(A) sequence. pTK305 (4112bp) [31] and pMC13 (3179bp) plasmids were used to produce (modified) mRNA containing respectively the luc2 or the eGFP reporter gene. These plasmids were constructed using standard Gateway cloning procedures and contain, besides the reporter genes also VEEV-derived 5′ and 3′ UTR.

*E. coli* bacteria containing the plasmids were cultivated in lysogeny broth (LB; Invitrogen, Massachusetts, USA) and the plasmids were subsequently isolated using the EndoFree Plasmid Maxi kit (Qiagen) when pDNA was used for injection or the Plasmid *Plus* midi kit (Qiagen) when further processed to mRNA.

### 2.3 mRNA synthesis

Template DNA was generated by linearizing the above plasmids using I-SceI endonuclease (NEB, Massachusetts, USA) before IVT with the MEGAscript T7 Transcription kit (Life Technologies, Massachusetts, USA). Post-transcriptional modifications were applied using the ScriptCap m7G Capping System, 2’-O-Methyltransferase kit and the A-Plus Poly(A) Polymerase Tailing kit (CELLSCRIPT, Wisconsin, USA). All mRNAs were purified with the RNeasy Mini kit (Qiagen) after IVT and after each modification. For production of modified mRNA, the uridine nucleotide was completely replaced by N1-methylpseudouridine (Tebu-bio, Belgium) during IVT. Correct translation of eGFP-encoding IVT-mRNAs and pDNA was verified after transfection of BHK cells using Lipofectamine MessengerMAX (ThermoFisher Scientific, Massachusetts, USA) and subsequent evaluation using the Nikon Eclipse Ti-S fluorescent microscope (Nikon, Belgium).

### 2.4 Injection, electroporation and lipid nanoparticles

Different doses of pDNA and mRNA (ranging from 0.01 to 10μg) were dissolved in 50μl PBS and 1 unit RNasin Plus RNase inhibitor (Promega) per μl solution was added to the mRNA before storing the solution at −80°C. Mice were sedated using the inhalation anesthetic isoflurane: 5% induction and 2% maintenance and shaved to allow for better exit of the photons produced during the luciferase-mediated conversion of luciferin. Injections were performed intradermally on one or both flanks of the mice using a 29G insulin needle (VWR, The Netherlands). Electroporation, when used, was executed immediately after each injection with a 2-needle array electrode containing 4 needles per row of 4 mm (AgilePulse, BTX Harvard Apparatus, Massachusetts, USA) as previously described [32]. In brief, 2 short high-voltage pulses were given (450V, 0.05ms) followed by 8 long low-voltage pulses (100V, 10ms) and an interval of 300ms between each pulse. Sa-RNAs coding for eGFP or luciferase were also encapsulated in LNPs like previously described [33]. Briefly, an ethanolic lipid solution was rapidly mixed with an aqueous solution containing sa-RNA at pH 4. The lipid solution consists of ionizable lipids, phosphatidylcholine, cholesterol and PEG in a 50/10/38.5/1.5 ratio, respectively. The RNA-loaded particles were characterized and subsequently stored at −80°C at a concentration of 1 μg/μl. The mean hydrodynamic diameter of these sa-RNA-LNPs was 72 ± 3 nm with a polydispersity index of 0.032 ± 0.009 and the zeta potential equaled −6 ± 1 mV (Nano ZS90, Malvern Pananalytical Ltd, Worcestershire, UK). The encapsulation efficiency was determined by RiboGreen RNA assay (as described before [34]) and equaled 95.8 ± 1.0%.

### 2.5 Bioluminescence imaging

Mice were intraperitoneally injected with 200μl D-luciferin (15 mg/ml, Gold Biotechnology, Missouri, USA) and *in vivo* bioluminescent imaging was performed 15 min later using an IVIS Lumina II (Perkin-Elmer, Zaventem, Belgium). The total flux in the region of interest was determined using the Living Image Software 4.3.1. *In vivo* bioluminescence imaging was repeated at different time points to study the expression kinetics of pDNA and the different mRNA vectors.

### 2.6 Statistical analysis

Statistical analysis and development of the graphs was performed with the software GraphPad Prism 6 (GraphPad Software, Inc., California, USA). Longitudinal experiments were analyzed with repeated-measures two-way ANOVA, followed by Sidak’s or Tukey’s multiple comparisons test. Fixed-time point analysis was performed using a ratio paired t-test and outliers were determined using Grubbs’ test. Differences are found to be significant when the p-value < 0.05 (*p < 0.05; **p < 0.01; ***p < 0.001; ****p < 0.0001).

## 3. Results

### 3.1 Electroporation increases the expression of sa-RNA and pDNA after intradermal injection

In a first set of experiments the expression kinetics after intradermal injection of pDNA or sa-RNA with or without subsequent electroporation was studied in BALB/c mice. The intradermal delivery route was selected as the skin is a large accessible organ containing many APCs for initiating an effective immune response. With both vectors, the luciferase expression could be detected as early as 5h after injection. Electroporation increased the expression of pDNA as well as sa-RNA. However, the beneficial effect of electroporation on pDNA transfection was only observed during the first five days (Fig. 1A). The peak in luciferase expression after intradermal electroporation of pDNA was reached after 2 days and at this time point electroporation induced a 3.5-fold increase in expression. The beneficial effect of electroporation was much more pronounced for sa-RNA (Fig. 1B). The luciferase expression profile of the sa-RNA was also different from that of pDNA. Sa-RNA reached its maximal expression 8-10 days after injection and at this moment, electroporation caused a 40-fold increase in expression compared to naked delivery. After 28 days the expression of the sa-RNA had declined to background, while the expression of pDNA lasted longer than 28 days. To allow for a better comparison of the expression levels, the areas under the curves (AUC) were calculated (Fig. 1C). After intradermal electroporation of sa-RNA the total amount of luciferase produced during the follow-up period was significant higher (30-fold higher) than after intradermal electroporation of pDNA (Fig. 1C). Furthermore, the overall expression of intradermal administered pDNA and sa-RNA was respectively 3-fold and 21-fold increased by electroporation. These data demonstrate again that especially the expression of sa-RNA and to a lesser extent pDNA is increased by electroporation and that the sa-RNA clearly outperforms pDNA after intradermal electroporation.

**Fig. 1.**
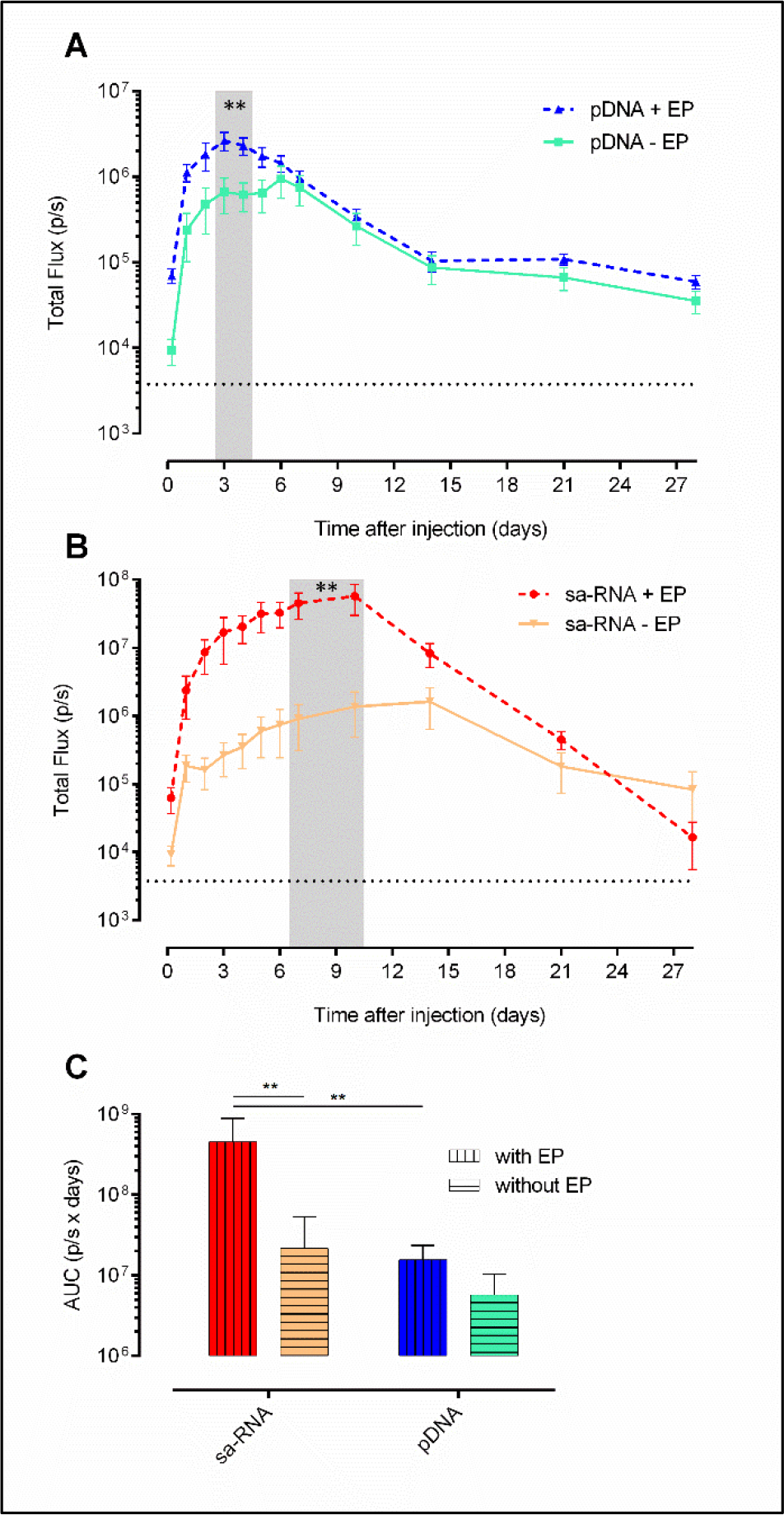
Effect of intradermal electroporation on the expression kinetics of sa-RNA and pDNA. The *in vivo* bioluminescence was measured at different time points after intradermal injection of 5μg of either pDNA (A) or sa-RNA (B), without (− EP) or with (+ EP) electroporation using needle array electrodes. Per group, three mice were injected on each flank (n=6). Results are shown as mean ± SEM of the total flux in the regions of interest (ROIs). The dotted black line represents the background signal. The shaded area indicates significant differences between the expression with and without electroporation. (C) AUC of the curves represented in A and B (mean ± SEM). **p < 0.01.

### 3.2 Comparison of the dose-dependent expression of sa-RNA, non-replicating mRNA and pDNA

Besides sa-RNA and pDNA, non-replicating mRNAs, whether or not containing modified nucleosides, are currently also intensively studied for vaccination, gene editing, or protein replacement therapy applications. However, a side-by-side comparison of the dose-expression profiles of these different synthetic mRNA platforms and pDNA after *in vivo* electroporation has not been performed. Therefore, we studied the luciferase expression levels and kinetics after intradermal electroporation of 10, 5 or 1μg sa-RNA, m1ψ-modified mRNA, unmodified mRNA or pDNA (Fig. 2A-C). The non-replicating mRNAs and pDNA reached, independent of the dose, their peak expression respectively 5h and 3 days after transfection. After this point the expression of the non-replicating mRNAs steadily decreased and disappeared at day 14 or day 10, when only 1μg was used. Plasmid pDNA showed after its peak a less steep drop and the expression of the two highest doses clearly remained above background till the end of the follow-up period. The expression profile of sa-RNA was different and reached, unrelated to the dose, a maximal plateau expression between day 3 and 10. After this plateau the expression showed a gradual drop and background levels were reached during the follow-up period (i.e. at day 28 and 21) with the two lowest doses (i.e. 5 and 1μg, respectively). The largest difference was reached 10 days after injection as sa-RNA was still inducing maximum luciferase expression while the non-replicating mRNAs had almost reached background level.

**Fig. 2.**
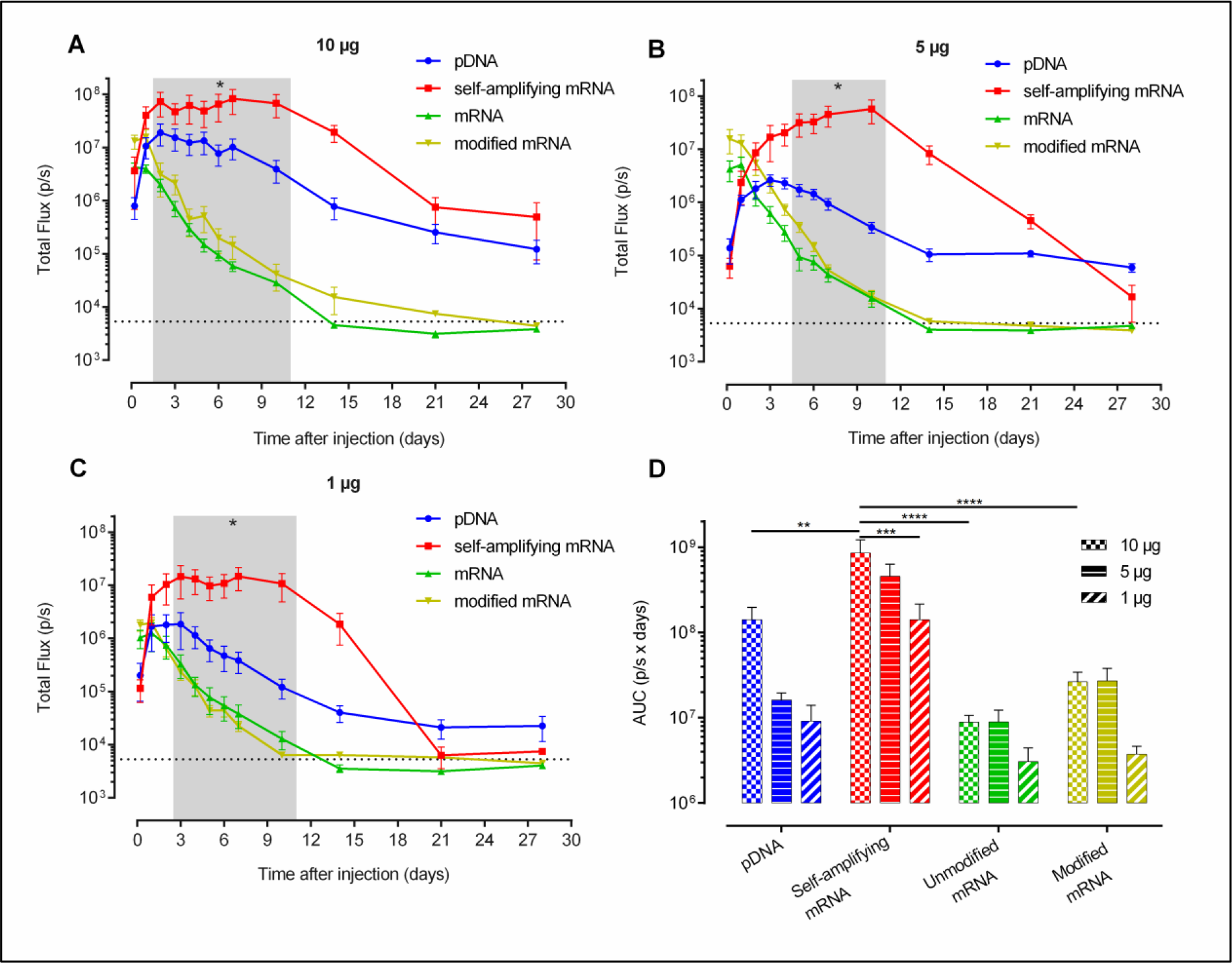
Dose-dependent expression kinetics after intradermal electroporation of pDNA, sa-RNA, unmodified and m1ψ-modified mRNA in mice. Three mice were intradermally electroporated in both flanks with 10μg (A), 5μg (B) or 1μg (C) of each vector (n=6). *In vivo* bioluminescence was measured over four weeks and mean ± SEM of the total flux in the ROIs is displayed. The dotted line represents the background signal. The shaded area indicates a significant difference between the luciferase expression of sa-RNA compared to pDNA, modified mRNA and unmodified mRNA (except for 10μg after 3 days (no significance) and 10μg and 1μg after 5 days (only significant difference between sa-RNA and the non-replicating mRNAs). (D) AUC (mean ± SEM) of the expression kinetic curves shown in A, B and C. *p < 0.05, **p < 0.01, ***p < 0.001, ****p < 0.0001.

To gain more insight into the dose-dependence, the total protein expression was evaluated by calculating the area under the curve (AUC) of the dose-response curves over a period of 4 weeks after injection (Fig. 2D). These AUCs nicely show a significant higher protein expression after sa-RNA transfection compared to pDNA and non-replicating mRNA. The m1ψ modification induced more protein expression compared to the non-modified variant (when 10 or 5μg was used), but less than pDNA or sa-RNA. A striking observation is that 10 or 5μg of the non-replicating mRNAs result in a similar expression curve and produce comparable amounts of luciferase during the follow-up period (Fig. 2A, B and D).

### 3.3 Sa-RNA at nanogram doses follows an all-or-nothing pattern

The high expression of 1μg of sa-RNA (Fig. 2C) and the fact that sa-RNA can amplify itself after intracellular delivery encouraged us to examine the expression when doses as low as 10ng were intradermally electroporated (Fig. 3). At these submicron doses the sa-RNA showed an all-or-nothing pattern: injection either resulted in a very high expression of luciferase or no expression at all. An overview of the success rates as a function of the sa-RNA dose is given in table 1. A high dose of 10μg was successful in 6 out of 6 attempts, while the lowest dose of 10ng only initiated expression in 1 out of 6 attempts. All successful injections resulted, independent of the dose (except at the lowest dose i.e. 10ng), in similar expression levels and curves with a maximal plateau expression between circa day 3 and 10. Submicron doses of pDNA were also assessed and resulted in much lower expression levels than sa-RNA and showed a clear dose-dependent profile. Ten ng of pDNA gave almost no luciferase expression whereas 10μg pDNA (Fig. 2A) reached peak luciferase levels comparable to those obtained with the successful injection of 10ng of sa-RNA.

**Table 1.**
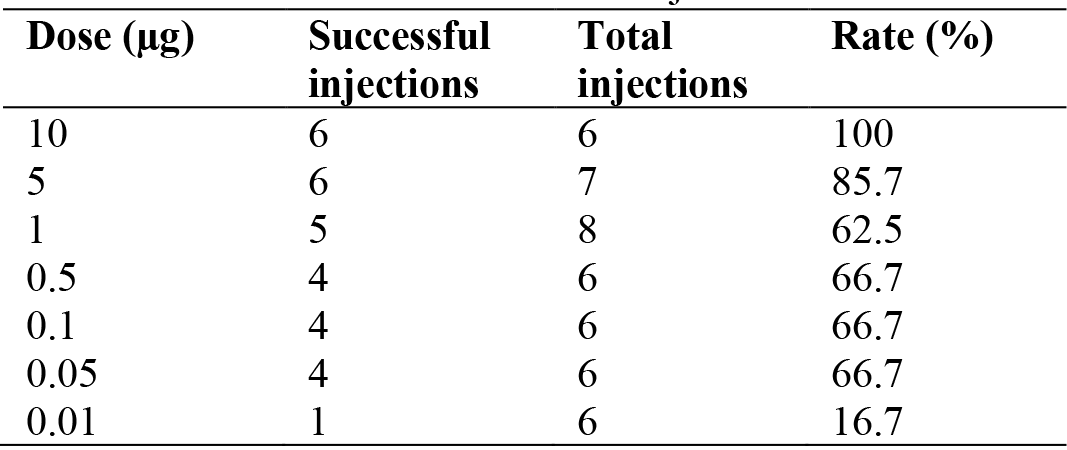
Rate of successful sa-RNA injections

| Dose (μg) | Successful injections | Total injections | Rate (%) |
| --- | --- | --- | --- |
| 10 | 6 | 6 | 100 |
| 5 | 6 | 7 | 85.7 |
| 1 | 5 | 8 | 62.5 |
| 0.5 | 4 | 6 | 66.7 |
| 0.1 | 4 | 6 | 66.7 |
| 0.05 | 4 | 6 | 66.7 |
| 0.01 | 1 | 6 | 16.7 |

**Fig. 3.**
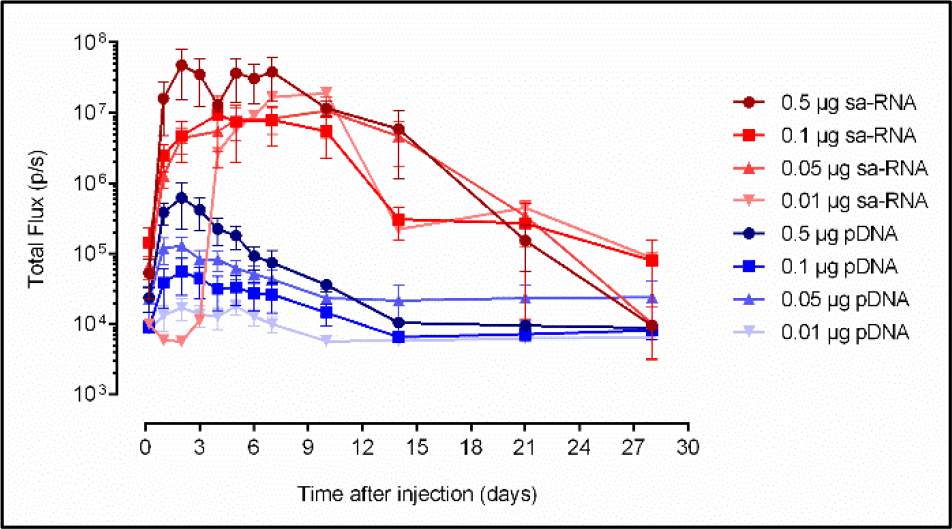
Expression kinetics of low doses of sa-RNA and pDNA after intradermal electroporation. Doses from 0.5 to 0.01μg of sa-RNA or pDNA were intradermally electroporated and luciferase expression was monitored during four weeks by *in vivo* bioluminescence imaging. Only sa-RNA administrations that resulted in a successful expression are used to calculate the mean. The expression success rates can be found in Table 1. In each group three mice were injected in each flank. Results are shown as mean ± SEM of the total flux in the ROIs.

### 3.4 Kinetics of the type I interferon response after intradermal electroporation of sa-RNA, non-replicating mRNAs and pDNA

To acquire information about the extent of the induced innate immune response, we evaluated the level and duration of IFN-β induction upon intradermal electroporation of the different synthetic mRNAs, pDNA and buffer in IFN-β reporter mice (Fig. 4). Electroporation of buffer did not provoke a notable increase in IFN-β, while all the vectors caused a clear IFN-β response that reached its maximum 8h after injection.

**Fig. 4.**
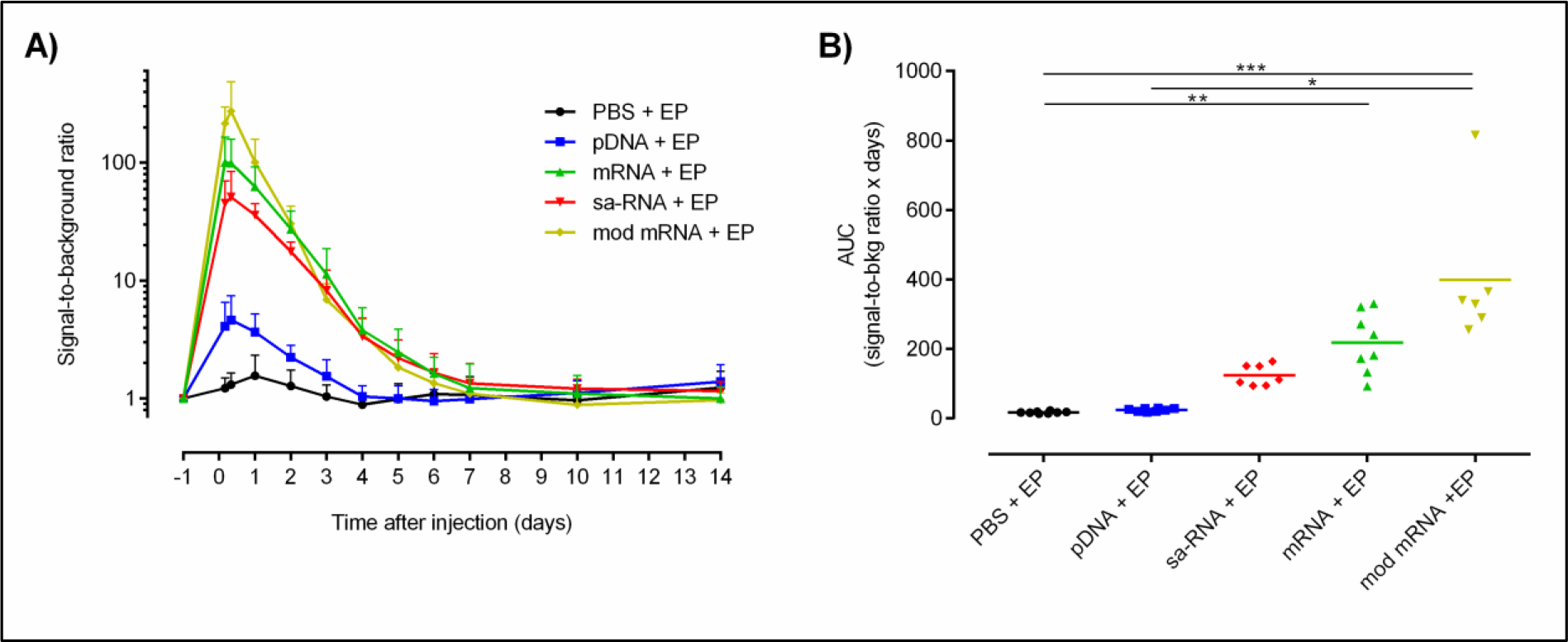
The extent and duration of the innate immune responses after intradermal electroporation of PBS, pDNA, sa-RNA, unmodified and m1ψ-modified mRNA. Five μg of each vector coding for eGFP was injected on the flank of heterozygous IFN-β luciferase reporter mice. (A) *In vivo* bioluminescence was measured starting 4h after injection. Signal to background ratio was calculated by dividing the bioluminescence signal in the ROIs (total flux, p/s) by its background signal at day −1 for each mouse. Mean + SD are displayed (n = 6-8). Significant differences are represented in Supp. Table 1. (B) As a measure of the overall IFN-β expression the AUCs of the curves in A were calculated. The horizontal bars represent the means. *p < 0.05; **p < 0.01; ***p

The highest IFN-β response was observed with the non-replicating mRNAs. Modified mRNA induced the highest average IFN-β induction shortly after electroporation, followed by unmodified mRNA. However, after 3 days, the IFN-β response induced by the mRNA’s becomes similar. IFN-β induction returns to baseline level 7 days after electroporation of mRNA. This led to an overall non-significant higher innate immune response induced by modified mRNA (Fig 4B). Self-amplifying mRNA caused, especially shortly after administration, a slightly lower IFN-β response than the non-replicating mRNAs. However, the IFN-β response dropped slower after administration of the sa-mRNA, resulting in an IFN-β response similar to that of unmodified mRNA from day 3 onwards. No significant induction of IFN-β was noticed after administration of plasmid DNA. The induction reached its maximum one day after administration and subsequently declined to background around day 4.

### 3.5 Electroporation versus lipid nanoparticle mediated delivery of sa-RNA

In a next experiment we compared the expression of 5μg sa-RNA formulated into state-of-the-art LNPs with that of naked sa-RNA with or without subsequent electroporation. Sa-RNAs formulated into LNPs showed completely different expression profiles compared to naked or electroporated sa-RNA after intradermal administration (Fig. 5A). LNP-mediated transfection of sa-RNA resulted in an early peak luciferase expression 24h after injection. After this time point the expression showed a very steep drop. However, after 7 days, a slight resurgence in luciferase expression occurred. As reported in Fig. 1, the expression of the naked sa-RNA was again clearly increased by electroporation. The shapes of the expression profiles of non-electroporated and electroporated sa-RNA were similar: a sharp increase during the first 24h and subsequently a steady increase until the maximal expression levels were reached around 8-10 days after injection. After day 10, the luciferase expression gradually dropped and 28 days after administration, the expression of the naked, electroporated and LNP delivered sa-RNA became close to background.

**Fig. 5.**
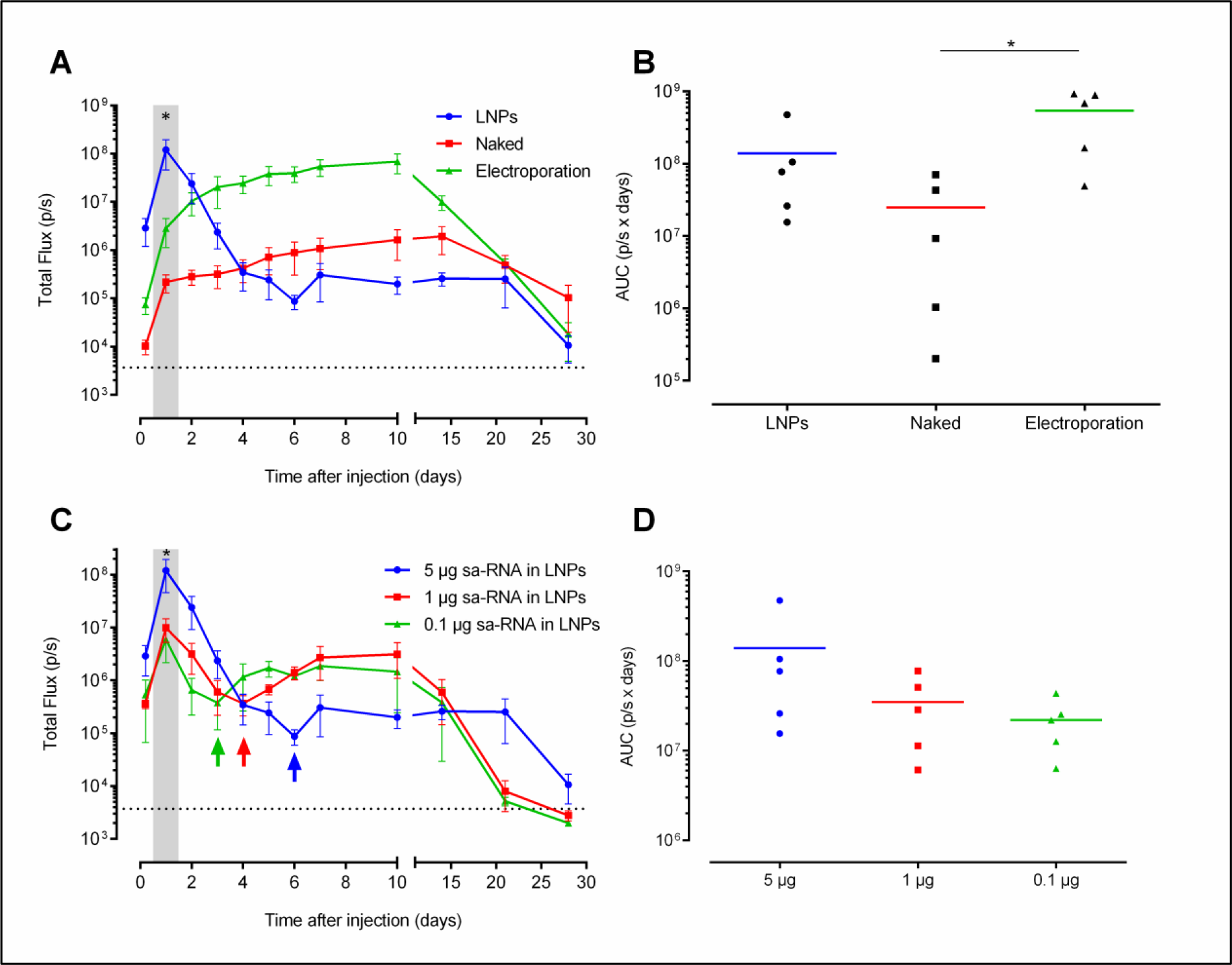
Lipid nanoparticle versus electroporation mediated intradermal delivery of sa-RNA. (A) Three mice per group were intradermally injected on both flanks with 5μg of either sa-RNA-LNPs, naked sa-RNA, or naked sa-RNA followed by electroporation. *In vivo* bioluminescence was measured over four weeks and mean ± SEM of the total flux in the ROIs is displayed. The dotted line represents the background signal and the shaded area indicates a significant difference between luciferase expression after use of LNPs versus electroporation. (B) AUC (mean ± SEM) of the expression kinetic curves shown in A was calculated and displayed to compare total protein expression between the different delivery systems. Individual values are plotted with the mean. (C) *In vivo* bioluminescence after intradermal injection of 5, 1 or 0.1μg of self-replicating mRNA encapsulated in LNPs. (D) The calculated AUCs of the curves in C. The shaded area indicates that the luciferase expression after injection of 5μg sa-RNA-LNPs is significantly higher than 1μg and 0.1μg. Mean ± SEM are displayed. The arrows indicate the time points at which the expression resurgences. *p < 0.05.

For certain applications such as protein (replacement) therapy the total amount of produced protein over time is more relevant than the peak expression level. Therefore, we calculated the area under the curves shown in Fig. 5A and C. The protein production after electroporation of the sa-RNA was respectively 22-fold and 4-fold higher than the amount of protein produced after injection of respectively naked or LNP formulated sa-RNA (Fig. 5B).

We also studied the dose-dependent expression after intradermal injection of sa-RNA-LNPs. Doses lower than 5μg, i.e. 1 and 0.1μg, of the LNP formulated sa-RNA resulted in similar but lower expression profiles (Fig. 5C, D). With the lower doses (i.e. 0.1 and 1μg) a similar resurgence of the luciferase is noticed as with the 5μg dose. However, this resurgence was more pronounced and the lower the dose the earlier this resurgence was noticed. All injections of sa-RNA-LNPs resulted in a successful expression.

### 3.6 Sa-RNA formulated in lipid nanoparticles cause expression in the draining lymph node after intradermal injection

During evaluation of the LNP-formulated sa-RNA, we observed a second, clearly delineated, smaller bioluminescent spot next to the injection spot (Fig. 6). This spot was located at the position where the subiliac lymph node can be found and hence suggested expression of the sa-RNA in the draining lymph node [35]. This additional expression spot could be seen until 48h after administration (data not shown).

**Fig. 6.**
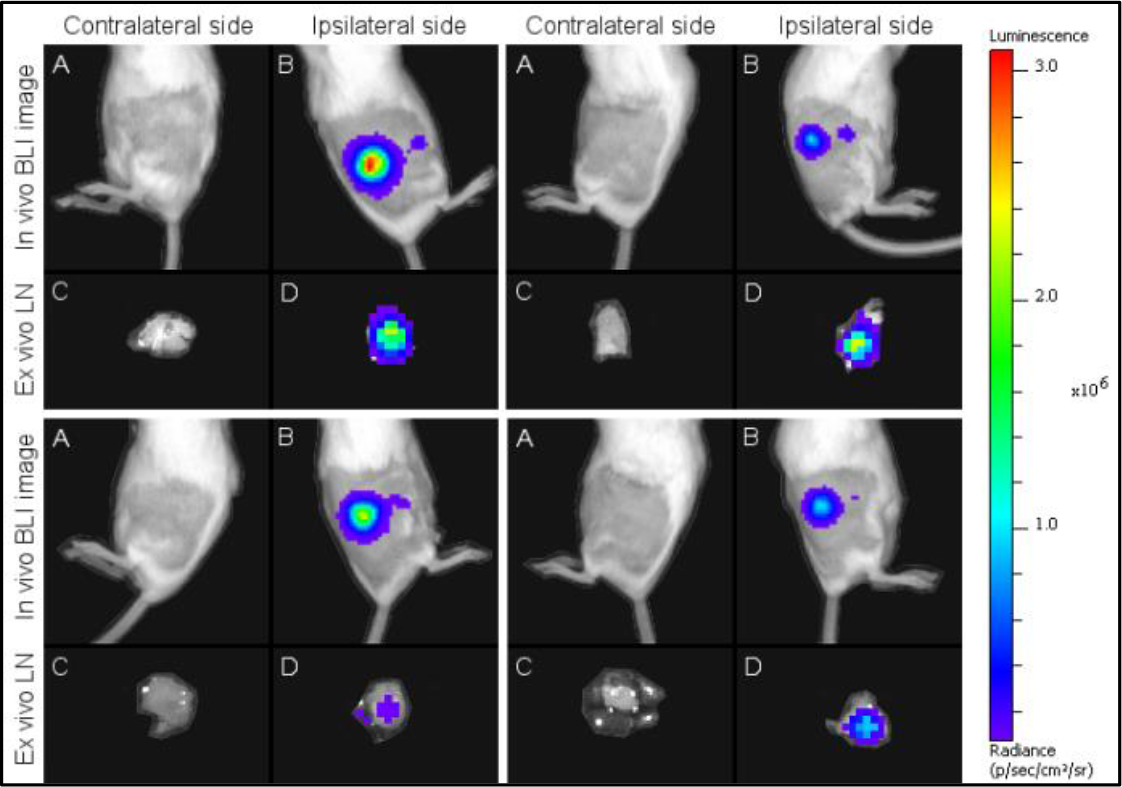
Bioluminescence images of four mouse after intradermal injection of 5μg self-replicating mRNA encapsulated in LNPs. Panels A and B show respectively *in vivo* bioluminescence images of the untreated contralateral (negative) flanks and treated ipsilateral (positive) flanks 5h after injection of sa-RNA-LNPs. Next to the bright bioluminescent spot, which is the injection spot, a small bioluminescent spot is visible at the location of the draining subiliac lymph node (panels B). Twenty four hours after injection of the sa-RNA-LNPs the subiliac lymph nodes were excised and *ex vivo* bioluminescent images were taken. Panels C and D show respectively the draining subiliac lymph nodes of the untreated contralateral (negative) flanks and treated ipsilateral (positive) flanks that received sa-RNA-LNPs.

To confirm this observation, mice were euthanized 24h after intradermal injection of sa-RNA-LNPs and the ipsilateral and contralateral (negative) subiliac lymph nodes were excised to be imaged *ex vivo* (Fig. 6 and Fig. 7). The ipsilateral (positive) subiliac lymph nodes showed significantly higher luciferase expression compared to the contralateral (negative) lymph nodes. This indicates that either transfected APCs travelled from the injection spot to the draining ipsilateral subiliac lymph node or that the sa-RNA-LNPs travelled to the ipsilateral subiliac lymph node and transfected there APCs and/or lymph node stromal cells. Such transfection in the draining lymph node was not observed after injection of the same dose of naked sa-RNA with or without electroporation.

**Fig. 7.**
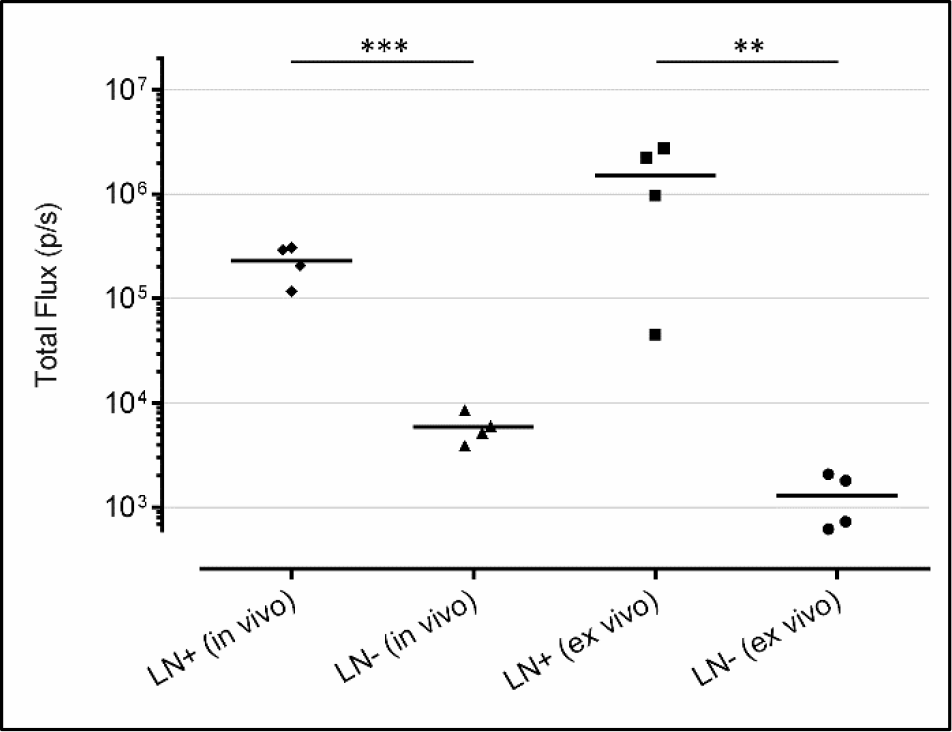
Comparison of the *in vivo* and *ex vivo* bioluminescence signals in the draining lymph nodes after intradermal injection of 5μg sa-RNA-LNPs. The *in vivo* bioluminescence signals were measured 5 h after intradermal injection. Twenty four hours after injection of the sa-RNA-LNPs mice were euthanized, the subiliac lymph nodes were excised and dripped with luciferin immediately before measurement. LN+= ipsilateral lymph node draining the injection spot. LN− = the contralateral lymph node. Horizontal bars represent the mean (n = 4). **p < 0.01; ***p < 0.001.

### 3.7 Kinetics of the type I interferon immune response after lipid nanoparticle and electroporation mediated delivery of sa-RNA

Self-amplifying mRNA formulated in LNPs will enter the cells through endosomes, which will likely lead to stimulation of the endosomal TLRs. It is expected that these TLRs get shunned when electroporation is used. To study whether this difference in cellular uptake affects the IFN-β response, we compared the kinetics of the IFN-β response after LNP and electroporation-mediated delivery of the sa-RNA (Fig. 8). Intradermal injection of 5μg sa-RNA-LNPs caused, compared to intradermal electroporation, a higher IFN-β response that also lasted longer: after LNP-mediated delivery of sa-RNA, IFN-β induction persisted for 14 days, while electroporation-mediated delivery only induced an IFN-β response during 7 days after injection. During the 14 day follow-up period the IFN-β response was about 8-fold higher after LNP mediated delivery.

**Fig. 8.**
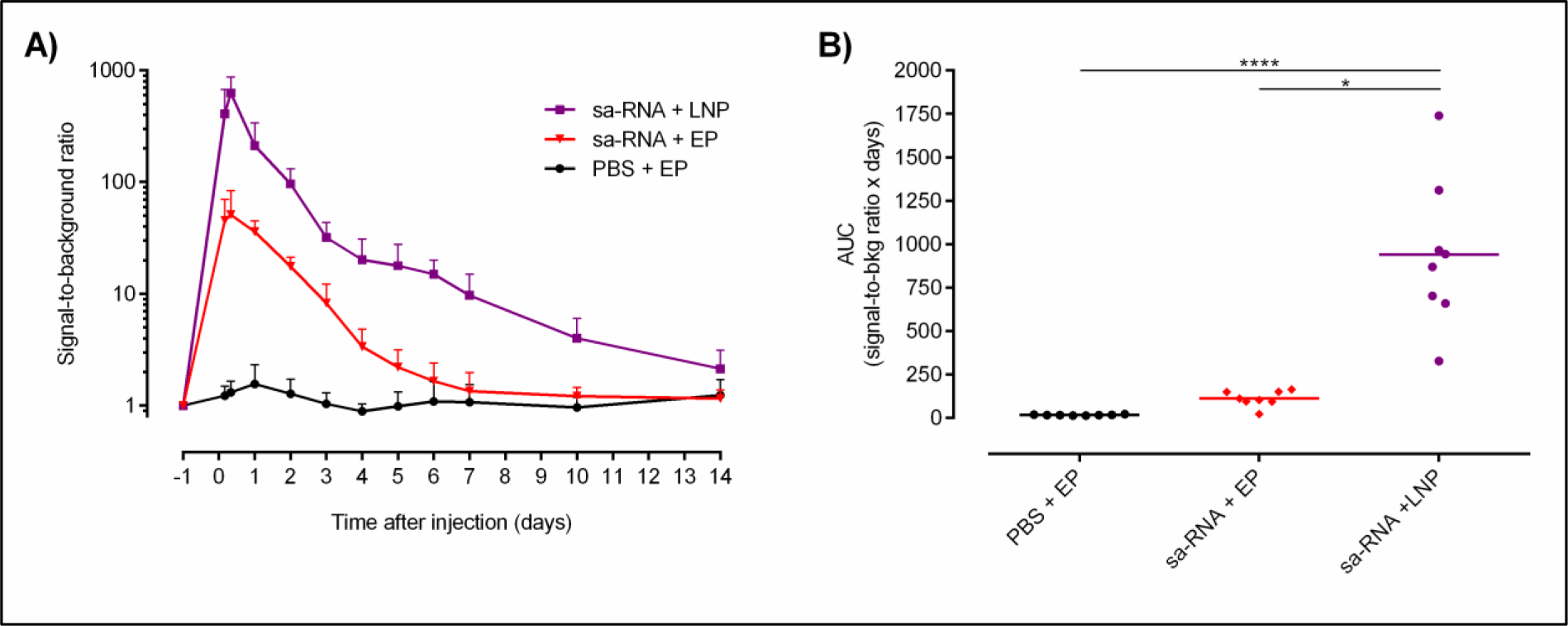
Kinetics of the type I interferon immune response after lipid nanoparticle mediated delivery of sa-RNA. (A) The extent and kinetics of the IFN-β induction in reporter mice after LNP and electroporation mediated intradermal delivery of 5μg sa-RNA encoding eGFP was measured by *in vivo* bioluminescence imaging. The *in vivo* bioluminescence was measured starting 4h after injection and was further monitored for 2 weeks. Signal-to-background ratios were calculated by dividing the bioluminescence signals in the ROIs (total flux, p/s) with its background signal at day −1 for each mouse. Mean ± SD are displayed (n = 8). Significant differences are represented in Supp. Table 1. (B) As a measure of the overall IFN-β expression the AUCs of the curves in A were calculated. Individual values are plotted with the mean. *p < 0.05; ****p < 0.0001

## 4. Discussion

The use of synthetic mRNA for the *in vivo* production of therapeutic proteins, like e.g. antibodies, erythropoietin or blood clotting factors, is recently gaining more and more attention [36–38]. For these applications a long-acting mRNA that can produce therapeutic proteins during several weeks would be ideal. Additionally, activation of the innate immune system by the delivered synthetic mRNA should be as low as possible or at least it should not affect the translation of the synthetic mRNA. Our data demonstrate that *in vivo* electroporation of a VEEV-based sa-RNA meets the above criteria. Electroporation of this sa-RNA caused a high and stable expression over two weeks. In contrast, non-replicating unmodified and modified mRNAs resulted in a much lower and shorter protein production. Sa-RNAs based on e.g. VEEV have been evaluated in the past as a possible new vaccination platform [6, 39–42]. However, the *in vivo* expression kinetics and especially the inherent innate immunity of VEEV-based sa-RNAs have not been studied in detail. We found that the sa-RNA caused, despite its intracellular amplification, a lower innate immune response than the non-replicating unmodified and modified mRNAs. Nevertheless, the kinetics of the IFN-β response was similar between sa-RNA and non-replicating mRNAs, with a maximal IFN-β induction 5h after administration followed by a steady decline. This may indicate that mainly the introduced sa-RNA, and not the replication of the sa-RNA, is causing the innate immune response. The absence of a strong and prolonged innate immune response after *in vivo* delivery of our VEEV-based sa-RNA can be explained by the capacity of the nsPs of VEEV to impair the signaling downstream of the type I IFN receptor [28]. Additionally, it has been reported that nsP2 can shut-off the translation of host mRNA by blocking its nuclear export [43–45]. These actions of the nsPs of VEEV may temper the innate immune response and prevent that it goes in full swing during the replication of our VEEV-based sa-RNA.

Sa-RNAs generate a replicase complex that is composed of the nsP1-4 of a single-stranded RNA virus. This raises the concern that these viral proteins may elicit an adaptive immune response, which would impede repeated administration of sa-RNAs. However, we have strong indications that such neutralizing adaptive immune response is not established. Indeed, as we observed that re-administration of our VEEV-based sa-RNA after four weeks did not result in a lower protein expression (unpublished data).

We showed that sa–RNAs are very potent vectors as they can cause gene expression at doses as low as 10ng after local administration in mice. However, it is important to mention that electroporation of 10ng of sa-RNA was only successful in 1 out of 6 injections. By evaluating the success rate as a function of the dose we found that the percentage of successful injections gradually dropped when the dose of sa-RNA was lowered. This observation may be attributed to the presence of ribonucleases (RNases) at the injection spot. RNases are abundantly present on the skin [46] and these RNases can contaminate the tip of the needle during injection. In this way minuscule amounts of RNases may be introduced and especially low doses of sa-RNAs will be very sensitive to such contaminating RNases.

To overcome activation of the RNA sensors, the mRNA can be rendered less immunogenic through incorporation of modified nucleotides like pseudouridine, 5-methylcytidine and N1-methylpseudouridine [31, 47–49]. However, incorporation of N1-methylpseudouridine-modified nucleosides in our non-replicating mRNA did not reduce the innate immune response and hence no significant increases in protein production were observed after electroporation of nucleoside-modified mRNAs. Similar results have been reported by Kauffman *et al.* who found that incorporation of pseudouridine modified nucleosides had no significant effect on either immunogenicity or protein expression of mRNA-LNPs after systemic injection [50]. In our work and in that of Kauffman *et al.* the mRNA was not purified by HPLC and hence double stranded mRNA or short aborted mRNA species that are known to be highly immunogenic were probably not completely removed, causing the innate immune response.

Although the expression of pDNA is lower than that of sa-RNA during the first weeks, pDNA is still an interesting vector for protein (replacement) therapy as its expression lasts longer than sa-RNA (up to 5 months after intradermal injection [51]). Another advantage is that the innate immune response after pDNA administration is very low and short-lived (Fig. 4A). However, the use of pDNA has some drawbacks like e.g. the theoretical risk of genomic integration and oncogenic mutagenesis, the presence of antibiotic resistance genes and the fact that for certain therapeutic proteins an uncontrolled expression during several months is not warranted.

To study the effect of different delivery methods on the *in vivo* performance of our VEEV-based sa-RNA, we encapsulated the sa-RNA into LNPs. Intradermal injection of these sa-RNA-LNPs resulted in a completely different expression profile than obtained after intradermal electroporation. Indeed, the expression of LNP formulated sa-RNA peaked shortly (i.e. 24h) after their administration. After this peak, the expression dropped sharply. Formulation of sa-RNAs into LNPs resulted thus in a faster and initial higher expression than obtained after electroporation mediated delivery. This indicates that LNPs cause a much more efficient intracellular delivery of the sa-RNA than electroporation (Fig. 5A). LNPs escort all mRNA into the cells without (excessive) degradation by extracellular RNases, while injection of naked mRNA in combination with electroporation presumably suffers from mRNA loss due to RNase degradation. Furthermore, electroporation does not direct all mRNA into the cells, resulting in a lower delivery efficiency. However, a strong innate immune response is induced as a consequence of the massive intracellular delivery of LNP formulated sa-RNAs (Fig. 8). This strong IFN-β induction is most likely responsible for the sharp drop in expression 24h after administration of the sa-RNA-LNPs. However, at day 3-6 the drop in expression stopped and a resurgence of the expression was noticed. We speculate that this can be linked to the observed kinetics of the innate immune response and the production of the innate immune evading nsP2 of VEEV. After delivery of the sa-RNA-LNPs, IFN-β immediately peaks and subsequently shows a gradual drop (Fig. 8A). At day 6 (when 5μg sa-RNA is used), the innate immune response drops below a threshold level (‘mRNA translation blockage threshold’), lifting the mRNA translation and allowing the persisting sa-RNA and subgenomic mRNAs to establish a revival of the expression. This resurgence occurred faster and was more pronounced when lower doses of sa-RNA-LNPs were used (Fig. 5C). Lower doses of the sa-RNA-LNPs are expected to induce a lower innate immune response. Hence, it takes less time for the innate immune response to drop below the “mRNA translation blockage threshold” and also more RNAs are expected to survive this shorter and less intense innate immune response. A similar resurgence, however, was not observed after electroporation of the sa-RNA. Therefore, the IFN-β response induced by electroporation of sa-RNA is most likely not strong enough to cause a decrease in luciferase expression.

A very interesting observation was the presence of a short-lived expression (up to 48h after administration, data not shown) in the draining subiliac lymph node after intradermal delivery of sa-RNA-LNPs (Fig. 6, 7). Expression of luciferase in the lymph nodes can be the result of the transport of the sa-RNA-LNPs towards the draining lymph nodes, or due to migration of transfected immune cells at the injection site towards the draining lymph nodes. Nevertheless, this expression in the lymph nodes most likely indicates transfection of antigen presenting cells. Therefore, formulation of sa-RNAs into LNPs is expected to genuinely boost the efficacy of sa-RNA vaccines as was also shown by Geall *et al.* [52, 53].

In conclusion, we demonstrated that *in vivo* intradermal electroporation of a VEEV-based sa-RNA outperformed the expression obtained after electroporation of pDNA or non-replication mRNAs. Furthermore, *in vivo* electroporation of our sa-RNA resulted in a short-lived and moderate innate immune response that did not affect the expression of the sa-RNA. When the VEEV-based sa-RNA was encapsulated in LNPs, a completely different expression and innate immune response profile was obtained. The expression rapidly peaked 24h after intradermal injection of sa-RNA-LNPs and subsequently showed a sharp drop which can be attributed to a massive induction of the innate immune system. Interestingly, intradermal injection of sa-RNA-LNPs also resulted in a protein expression in the lymph nodes, which supports the potential use of sa-RNA-LNPs as vaccines. However, it needs to be examined whether the induced innate immune response after administration of the sa-RNA-LNPs is balanced enough to potentiate adaptive immune responses. Indeed, it has been shown that a too high innate immune response can also be detrimental for the adaptive immune response [54, 55].

## 5. Acknowledgements

We would like to thank Tasuku Kitada and Ron Weiss of the Massachusetts Institute of Technology for kindly providing us with the pTK160 and pTK305 plasmids. This research has also benefitted from a statistical consult with Ghent University FIRE (Fostering Innovative Research based on Evidence).

